# Multimodal Thrombectomy Device for Treatment of Acute Deep Venous Thrombosis

**DOI:** 10.1101/2022.02.21.481340

**Authors:** Usama Ismail, Roger A. Rowe, John Cashin, Guy M. Genin, Mohamed Zayed

**Author notes:** These authors contributed equally to the work (Guy M. Genin, Mohamed Zayed).

## Abstract

Deep vein thrombosis (DVT) is a potentially deadly medical condition that is costly to treat and impacts thousands of Americans every year. DVT is characterized by the formation of blood clots within the deep venous system of the body. If a DVT dislodges it can lead to venous thromboembolism (VTE) and pulmonary embolism (PE), both of which can lead to significant morbidity or death. Current treatment options for DVT are limited in both effectiveness and safety, in part because the treatment of the DVT cannot be confined to a defined sequestered treatment zone. We therefore developed and tested a thrombectomy device that enables the sequesteration of a DVT to a defined treatment zone during fragmentation and evacuation. We observed that, compared to a predicate thrombectomy device, the sequestered approach reduced distal DVT embolization during *ex vivo* thrombectomy. The sequestered approach also facilitated isovolumetric infusion and suction that enabled clearance of the sequestered treatment zone without significantly impacting vein wall diameter. Results suggest that our novel device using sequestered therapy holds promise for the treatment of high risk large-volume DVTs.

## Introduction

Venous thromboembolism (VTE) is the third most common vascular disease in the world^1^, and leads to an estimated 60,000-100,000 annual deaths in the US annually^2^. Deep vein thrombosis (DVT) is one of the major causes of VTE, and occurs when abnormal blood clots (thrombus) develop within the venous system^2^. Venous thrombus can disassociate and travel into the pulmonary arteries, causing a potentially fatal pulmonary embolism (PE)^3^. Individuals with large-volume DVTs in the proximal veins of the pelvis and abdomen (in the iliac veins and vena cava) are at higher risk of suffering a PE^4^. Therefore, there is a pressing need for effective and safe treatment options for the management of higher risk large-volume DVTs.

Two widely used options exist for the treatment of patients with large-volume DVTs in the large veins of the abdomen and pelvis (Figure 1), but both have significant limitations. Both of these approaches involve endovascular insertion of catheter devices to perform thrombectomy (fragmentation and/or removal of DVT). The first of these treatment options utilizes percutaneous mechanical thrombolysis catheters. These aim to fragment and dissolve large-volume DVT within the venous lumen into smaller fragments (Figure 1, left panel). However, this fragmentation comes with the risk of damaging the inner vein wall during the mechanical sheering of thrombus^5, 6^. Additionally, the procedure is associated with high risk of distal embolization, in which smaller thrombus fragments travel with the venous blood flow to the heart and lungs causing a PE.

**Figure 1.**
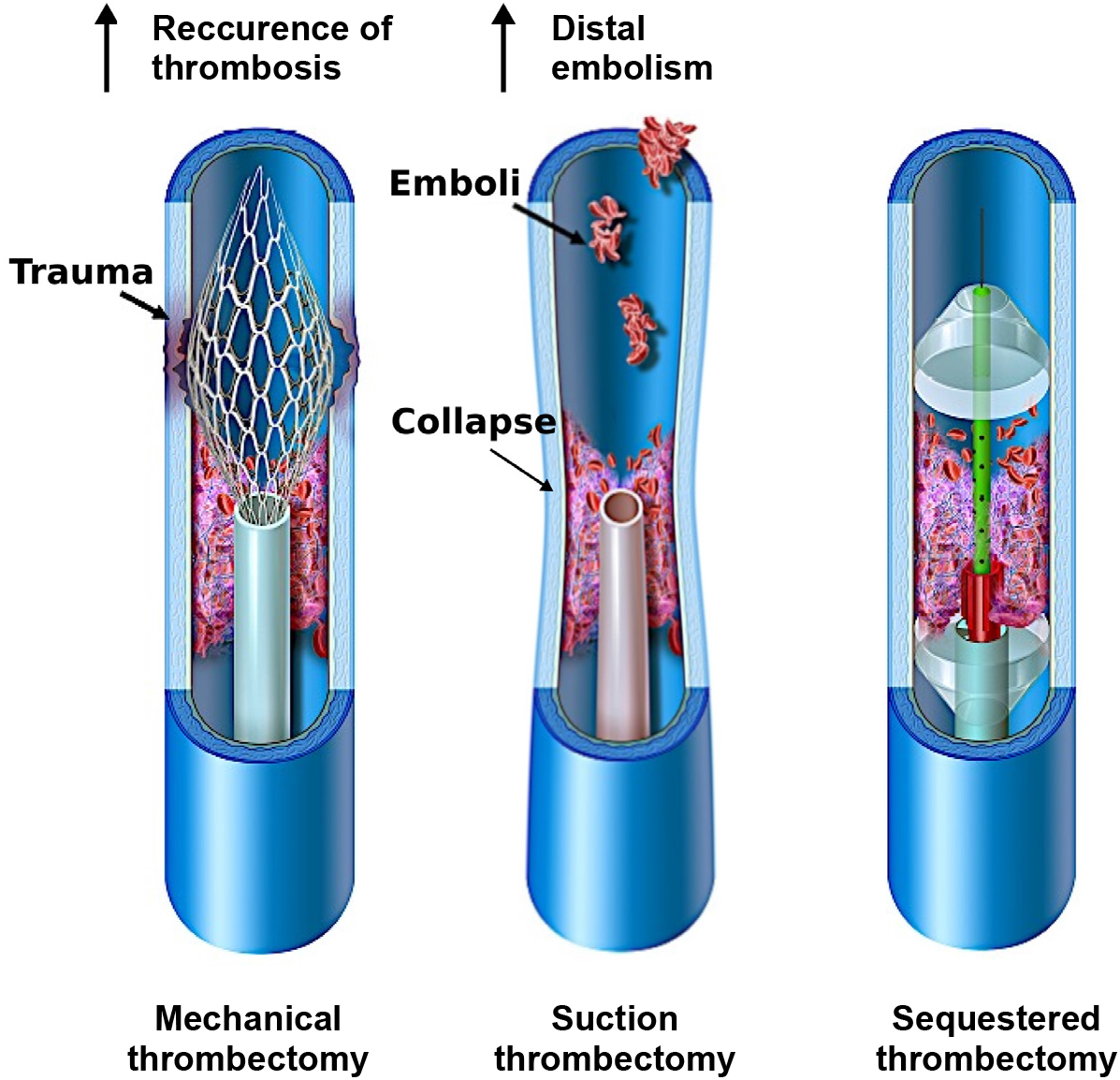
Comparison of traditional thrombectomy treatment options and a novel multi-modal sequestering thrombectomy catheter.The graphics in this figure was drawn for Caeli Vascular by Matt Holt

An alternative treatment option is suction thrombectomy (Figure 1, middle panel). Here, suction devices inserted through the vasculature are used to debulk large-volume DVT via negative pressure evacuation. However, suction comes with a risk of vein wall collapse, which can lead to trauma on the inner lumen of the vessel, as well as risk of distal embolization of thrombus fragments that separate from the main thrombus but are not successfully removed^7^. Additionally, substantial blood loss can occur during these procedures as thrombus fragments are removed along with variable quantities of blood volume.

Given the inherent limitations of existing mechanical and suction thrombectomy devices, we proposed a multi-modal design that sequesters the treatment zone (Figure 1, right panel). The aim of this novel device would be to reduce the risk of distal thrombus embolization during a thrombectomy procedure. Additionally, the device would allow sequestered flushing of thrombus fragments within the defined treatment zone without traumatically impacting the vein wall. We hypothesized that in an *ex vivo* setting, this novel sequestering thrombectomy device would effectively remove thrombus, reduce distal embolization, and prevent vein wall over-distension or collapse.

## Methods

### Prototype of a catheter with a sequestered treatment zone

A thrombectomy device that sequestered a treatment zone between two balloons was developed in collaboration with Nordson Medical, Inc. (Marlborough, MA). The multi-modal sequestered thrombectomy catheter (MMSTC), was made of multiple extruded tubes, with multiple inner lumens. The proximal, outermost catheter was 1 meter long, and had an outer diameter of 7.33 mm (22 French (Fr), where 1 Fr = 1/3 mm is the standard unit to measure diameters of catheters) and an inner diameter of 7 mm (21 Fr). The proximal catheter was made of polyether block amide (Pebax 5533, Arkema, Colombes, France), and coated to make the surfaces hydrophobic. This catheter was inserted over a guide wire (0.889 mm (0.035 inch) diameter stainless steel wire), and terminated in a proximal balloon that could be inflated to demarcate and seal the the proximal end of the treatment zone (left side, Fig. 2A).

**Figure 2.**
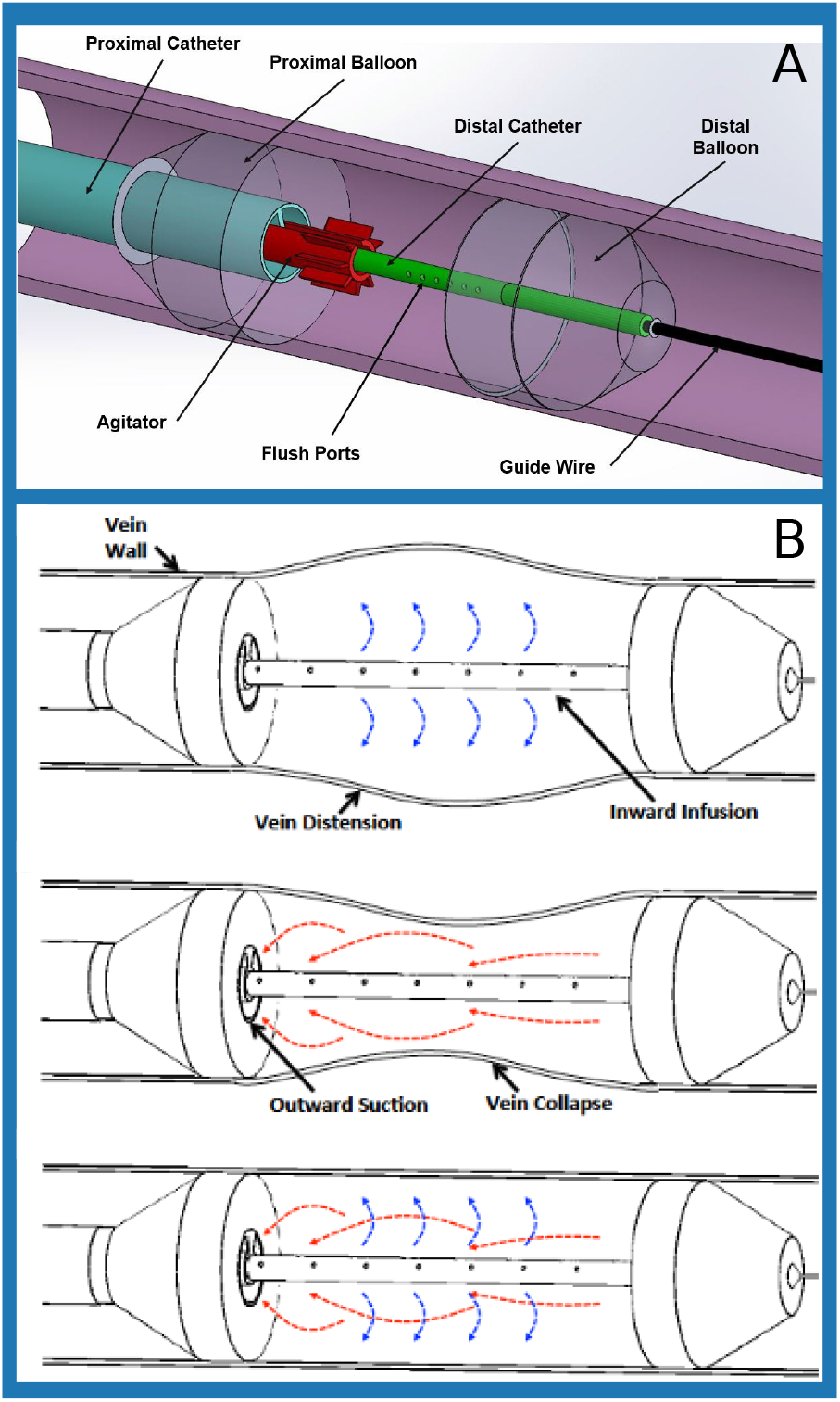
A) Components of the multi-modal sequestering thrombectomy catheter that isolates the treatment zone.B) Comparison of infusion only, suction only, and infusion and suction modes of catheter operation to evacuate thrombus fragments within the isolated treatment zone.

The second component of this device was a thinner but longer distal catheter that could be inserted over the same guide wire through through the lumen of the proximal catheter (right side, Fig. 2A).. This terminated in a second, distal balloon that defied the distal end of the treatment zone. Inside distal catheter was 1.1 m long and had a 2.33 mm (7 Fr) outer diameter. This multi-lumen extrusion was made of nylon 12 ([(CH_2_)_1_1C(O)NH]*_n_*). The distal catheter included flush ports that enable the surgeon to infuse drugs such as thrombolytics, and also to subsequently introduce fluid to the treatment zone that flushes fragments of thrombus down one of the lumens of the proximal catheter; simultaneous suction can be applied to that lumen.

The MMSTC included a center-line, rail-mounted mechanical agitator for the purpose of fragmenting thrombus, and aimed to do so while minimizing the risk of vein wall contact and trauma. The plastic agitator measured 5mm x 15mm and tracked over the outer surface of the distal catheter and that could be moved independently within the treatment zone to fragment thrombus and using 1mm stainless steel wire cannulas.

The user end of the device utilized standard luer-lock components from the Nordson Medical, Inc. catalog which were over-molded onto the thrombectomy device. The MMSTC thus integrated both mechanical and suction thrombectomy modalities to remove large-volume DVTs within a sequestered region of vascular segment. The catheter’s adjustable double-balloon system aimed to isolate and sequester the treatment zone to avoid distal thrombus embolization during treatment sessions. As described below, we tested the hypothesis that, in an *ex vivo* system with an artificially generated blood clot, the MMSTC catheter system would effectively remove thrombus fragments, reduce distal embolization, and prevent vein wall over-distension or collapse.

### Isovolumetric flushing of the sequestered treatment zone

The MMSTC catheter system facilitated isovolumetric infusion and suction through simultaneous infusion and suction of liquid in the sequestered treatment zone (Fig. 2B) with the aim of minimizing vein wall distension or collapse during evacuation of thrombus fragments. This was achieved via a linear actuator device to simultaneously infuse and suction fluid through the treatment at equal volumetric flow rates (Fig. 3).

**Figure 3.**
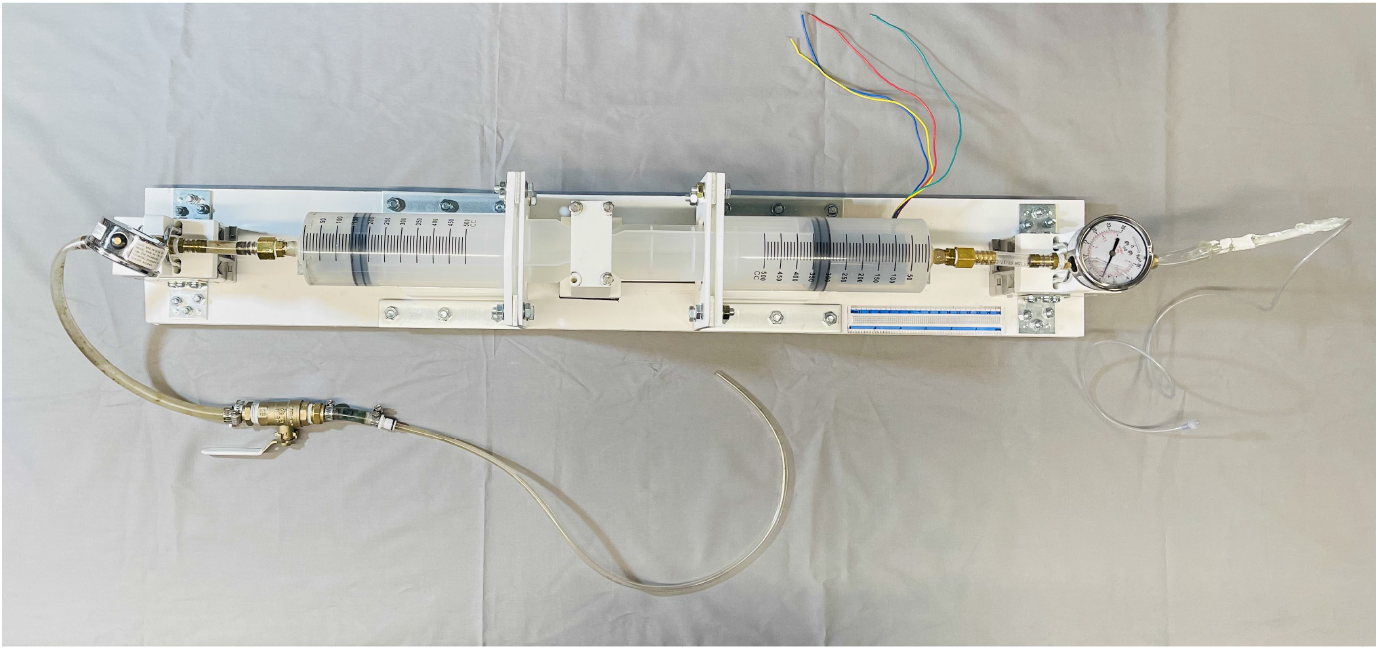
A top-down view of the linear actuator used in conjunction with the Hydra Catheter.

The linear actuator was composed of a motor-powered cart which followed a lead screw along the axis of the device and displaced a 3D printed coupler that connected to the the plungers of two 500 ml syringes. Movement of the coupler thus depressed the plunger of one syringe while retracting the plunger of the other syringe at the same rate. Surgical tubing connected the end of the positive pressure syringe to the to the infusion lumen of the distal catheter, and connected the end of the negative pressure syringe to the evacuation lumen of the proximal catheter. A pressure gauge was mounted between the positive pressure syringe and the infusion lumen.

A US National Electrical Manufacturers Association (NEMA) standard size 23 motor (faceplate 58 x 58 mm) was used to drive the cart, and was controlled by a microcontroller (Arduino Mega 2560 Rev3, Arduino, Ivrea, Italy), a Tb6560 motor driver (TB6560 3-axis stepper motor controller, Hailege, Shenzen, China), and a 24 V DC power supply (RS-150-24, Mean Well, Taipei, Taiwan). A range of motor speeds was tested.

### Thrombectomy testing in an ex vivo setting

The efficacy of the MMSTC and an industry-standard catheter (Indigo CAT 5 Fr, Penumbra, Alameda, CA) were assessed in an *ex vivo* circulation system that contained a model thrombus within a segment of 19.05 mm diameter clear vinyl tubing (3/4″ ID clear vinyl tubing, Abbott Rubber Company, Inc, Itastca, Illinois, USA). Both devices use a combination of suction and physical disruption of the thrombus to treat thrombus. Each was used to treat a section of in-lab made thrombus for 30 seconds, an interval of time that is clinically relevant for our own surgical practice, and device efficacy was assessed by the weight of thrombus collected after 30 seconds of treatment.

#### Model thrombus generation

Our model thrombus model for these experiments was created by mixing bovine blood, deionized water, and flour for a thickening agent. Sourcing bovine blood from a local meat packing plant (Star Packing Company, St. Louis, Missouri, USA), a 6.5 mL mixture of 6:4 blood to water was mixed together with one teaspoon of flour as a thickening agent. Next, this mixture was poured into a portion of flexible vinyl tubing and allowed to rest in a 4 degree Celsius refrigerator for 24 hours. The next day blood that had risen to the surface of the mixture was drained and another teaspoon of flour was mixed into the solution. The final mixture was allowed to rest at 4 degrees Celsius for another 24 hours.

#### Laboratory flow model

The testing setup used to assess catheter performance and the mass of distal emboli generated during a simulated surgical thrombectomy consisted of a reservoir of water that was pumped through a segment of model thrombus within a tube at a rate of 1500 ml/s using a chemical pump (3388 Variable-Flow Chemical Pump, Control Company, Webster, Texas, USA). The setup (Fig. 4) consisted of a main central channel housing the thrombus and treatment zone, with the model thrombus supported in by a 3D printed holder. A port allowing introduction of a catheter was included, as was an alternate overflow route representing peripheral vasculature that accommodates blood return in the case of venous blockage. This also helped prevent the introduction of air into the system. The treatment zone drained to an outlet hose that sent passed saline and distal emboli to a mesh sieve with a coffee filter (porosity of 10 – 20*μ*m, Simply Best™ 8-12 cup coffee filters, Topco Associates, LLC, Elk Grove Village, Illinois, USA) paper, for collection of distal emboli. In all experiments, the chemical pump was run and allowed to fill the flow setup tubing until no air bubbles were evident visually.

**Figure 4.**
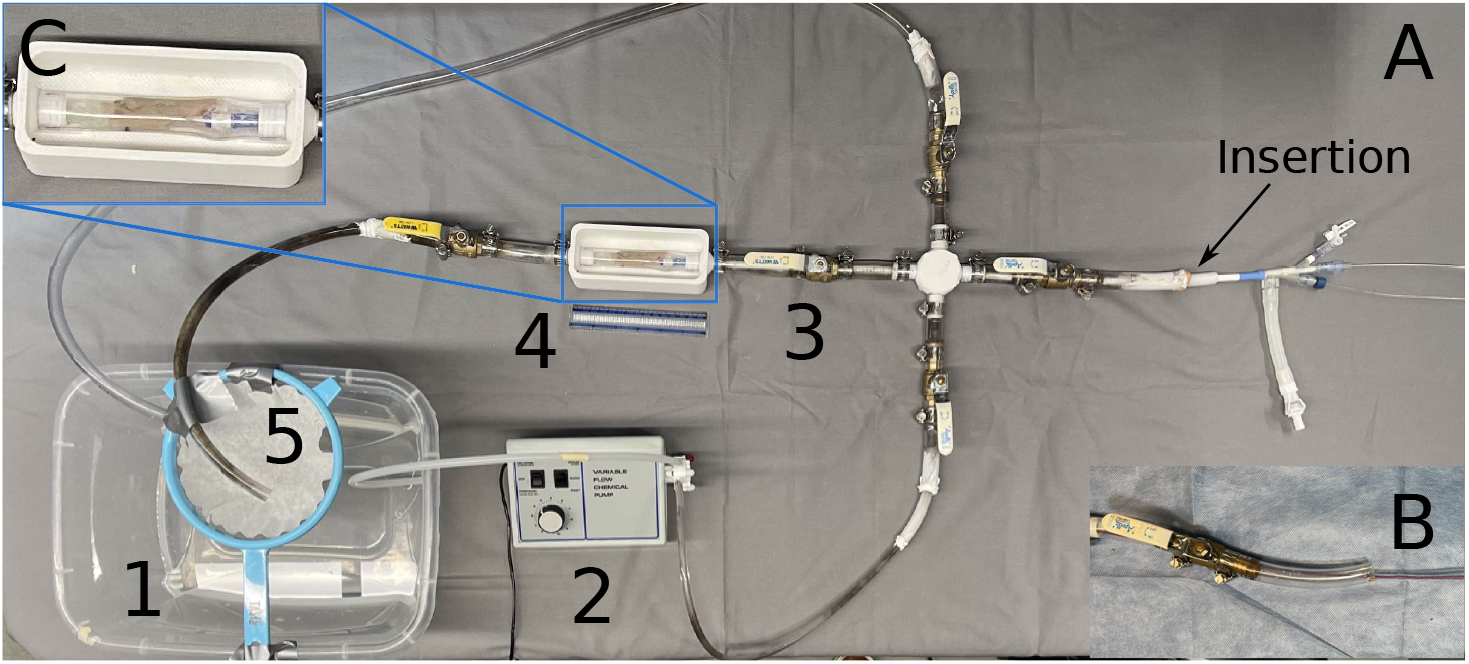
A) The laboratory flow model consisted of (counter-clockwise, beginning in the lower left) a saline reservoir (1); a pump (2); a valve that enabled introduction of a catheter and allowed for overflow (3); a visualization chamber that supported either a clear tube or a segment of porcine vena cava with a laboratory-made thrombus (4); and a filtered return that enabled collection of any embolized thrombus (5). The MMSTC device is shown entering the setup at the insertion point. B) The Penumbra device is shown entering the insertion point. C) A zoomed in view of the treatment area/thrombus model of the laboratory flow model.

#### Simulated surgical thrombectomy procedures

To test the MMSTC, the catheter was introduced through the catheter port (right side, Fig. 4), the proximal balloon was inserted and then inflated, and the distal catheter was then advanced through the model thrombus and its balloon inflated. An endoflator (BasixTouch™ Inflation Device, Merit Medical, South Jordan, Utah, USA) was used to inject a 1:1 mix of contrast agent and deionized when filling balloons. The linear actuator was used at a motor speed of 300 RPM to infuse liquid and suction thrombus while the agitator was displaced along the length of the treatment zone to disrupt the thrombus segment. After 30 seconds, the thrombus collected in the linear actuator was collected in a pre-weighed coffee filter and allowed to dry overnight (cf.^8^) to calculate the dry weight of the total thrombus collected.

To test the predicate Penumbra Indigo 5 Fr catheter, an Indigo CAT 5 Fr Separator (Penumbra, Alameda, CA) was used to fragment the thrombus, and a vacuum regulator, tubing, and 1000 ml collection canister set (Gomco Model 405 Table Aspirator, Allied Healthcare Products Inc, St. Louis, Missouri, USA) was used to generate suction and collect thrombus. The vacuum pump was set to 75 kPa (22.5 inHg) of pressure. The thrombus collected in the collection canister was collected on filter paper, dried overnight, and weighed.

### Quantification of distal embolization in situ

Both devices were tested to assess the amount of distal embolization generated during a simulated thrombectomy conducted in the flow circuit described above. A piece of tubing filled with the simulated thrombus described above was attached to the 3D printed housing in the flow circuit. A catheter was inserted into the flow setup as described above.

For the MMSTC, to test the efficacy of the sequestered approach to thrombectomy, the balloons were either left uninflated (control), inflated 50%, or inflated 100% as above. For the MMSTC, the agitator was used to fragment the thrombus, and the isovolumetric pump was engaged. While this treatment proceeded over 30 s, any thrombus that escaped the sequestered treatment zone was collected in the pre-weighed coffee filter and its dry weight recorded.

For the Penumbra Indigo 5 Fr device, the same setup was used, but as above, instead of the linear actuator, the Gomco Allied 406 Vacuum regulator was used at 85 kPa (25 inHg) to induce suction in the catheter, and a Penumbra separator (Indigo System CAT 5 Separator, Penumbra Inc., Alameda, CA) was used in place of the Hydra agitator to break up the thrombus.

### Measurement and prediction of vein wall distention and collapse ex vivo

To assess how the MMSTC system strained the vessel wall during isovolumetric evacuation of the sequestered treatment zone, the degree to which the process distorted the vein wall was monitored by tracking the hoop stretch ratio *λ_θ_* (*p*) in a bovine vena cava specimen acquired from a local red abattoir that was inserted in place of the model thrombus and clear tubing in the flow loop. The stretch ratio followed:

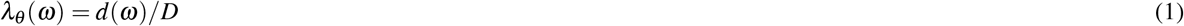

where *D* is the initial diameter of the vena cava, and *d*(*ω*) is the steady state diameter for running of the syringe driver motor at a rotational speed *ω*. Vena cava segments were stored at 4°C in 10% formalin, then tied to two connectors at the center of the flow loop using hose clamps. For these tests, the MMSTC was inserted into the vena cava segment and the balloons were inflated to 100%. Before the experiment, the original diameter, D, of the middle cross-section of the vena cava was measured using a set of calipers. After this, the linear actuator was operated in one of three ways. First, the rotational speed-stretch ratio relationship was measured only the infusion syringe connected to the MMSTC, but not the suction syringe (infusion only). Second, The relationship was measured with the suction syringe but not the infusion syringe connected to the MMSTC (suction only). Finally, the relationship was measured with both syringes connected (isovolumetric mode). While the linear actuator was running the diameter, *d*(*ω*), of the middle cross-section of the vena cava was taken again with the calipers. This process was repeated three times for each mode at 200, 300, and 400 motor RPM.

For comparison, two mathematical models for the mechanical behavior of a thin walled tube were studied. The first was a model of a hyperelastic tube under inflation, which, as described in the supplement, which predicts how inflation pressure, *p*, relates to *λ_θ_*:

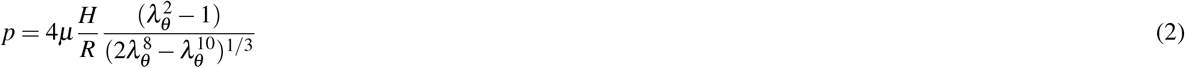

where *H* is the thickness of the vein prior to inflation, *R* is the radius of the vein prior to inflation, and *C*_1_ is the shear modulus of the vein.

The second was the classical Bryan solution for the buckling pressure *p_critical_* of a long tube that is clamped at its ends^9^:

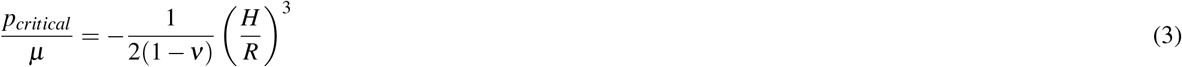

where *v* = 0.5 is Poisson’s ratio for the vein wall.

### Statistics

Differences between means was assessed using one-way ANOVA, with a 5% false positive rate taken as the threshold for statistical significance. All results were reported as mean ± standard error to the mean. Each experiment was repeated a minimum of three times.

## Results

### Thrombectomy efficiency

Both the MMSTC and the Penumbra Indigo CAT 5Fr catheter removed substantial thrombus when used for 30 seconds in our flow system to remove a model thrombus. The MMSTC removed significantly more thrombus over this time interval: the Penumbra Indigo CAT 5 Fr catheter removed 27± 3.14 mg, while the MMSTC removed 87± 3.86 mg (*p* = 0.005 using one-way ANOVA; Fig. 5).The flow model was a crude representation of a vascular surgical treatment zone since it does not have the variable shape and mechanics of a natural *in vivo* venous structure. The metric chosen for device performance in this *ex vivo* model was therefore not the fraction of thrombus removed, but rather the size distribution of thrombus particles collected in the distal filter spliced into the flow model. The latter was considered critical since successful multi-modal suction of the thrombus particles is highly dependent on thrombus fragmentation to avoid obstruction of the suction catheter by large thrombus pieces.

**Figure 5.**
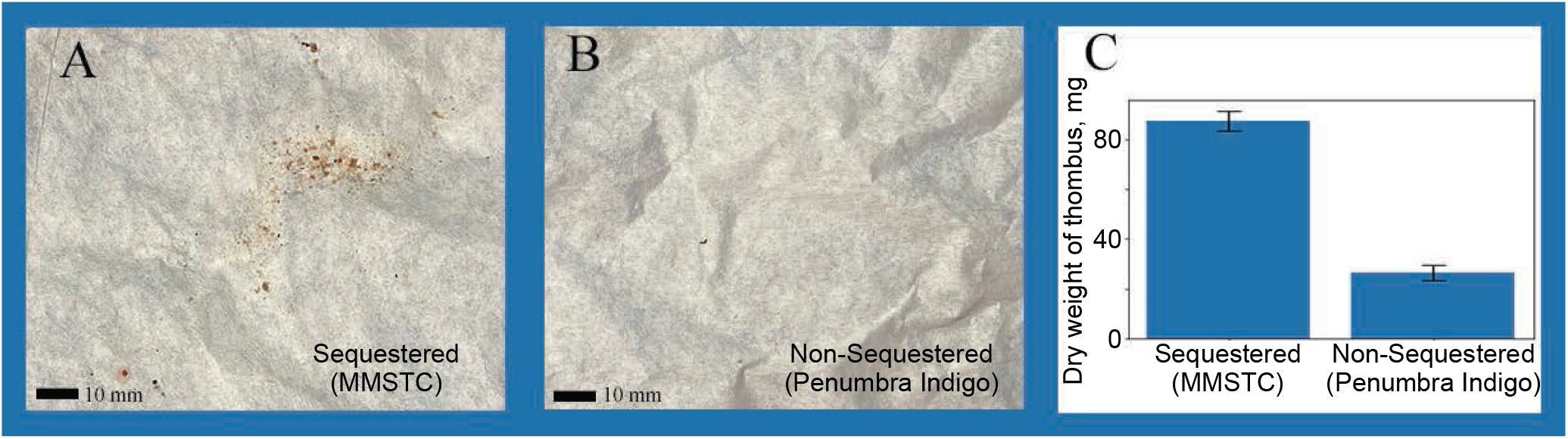
A) Thrombus evacuated by the MMSTC in a typical experiment, viewed on filter paper, over the course of a 30 second treatment within the flow system. B) Thrombus evacuated by the Penumbra Indigo CAT 5 Fr catheter in a typical experiment, viewed on filter paper, over the course of a 30 second treatment within the flow system. C) The MMSTC evacuated significantly more thrombus than the Penumbra Indigo CAT 5 Fr catheter under these conditions (*p* = 0.005).

### Sequestering the treatment zone decreased distal embolization of thrombus

Distal thrombus embolization was evaluated in both catheters. To test the hypothesis that the sequestering of the treatment zone reduced distal embolization, the MMSTC was operated with the balloons that sequester the treatment zone inflated to 0%, 50%, or 100% of the maximum inflation (Fig. 6A-C). We additionally compared the MMSTC to the Penumbra Indigo CAT 5 Fr catheter that does not have a distal embolization protection feature.

**Figure 6.**
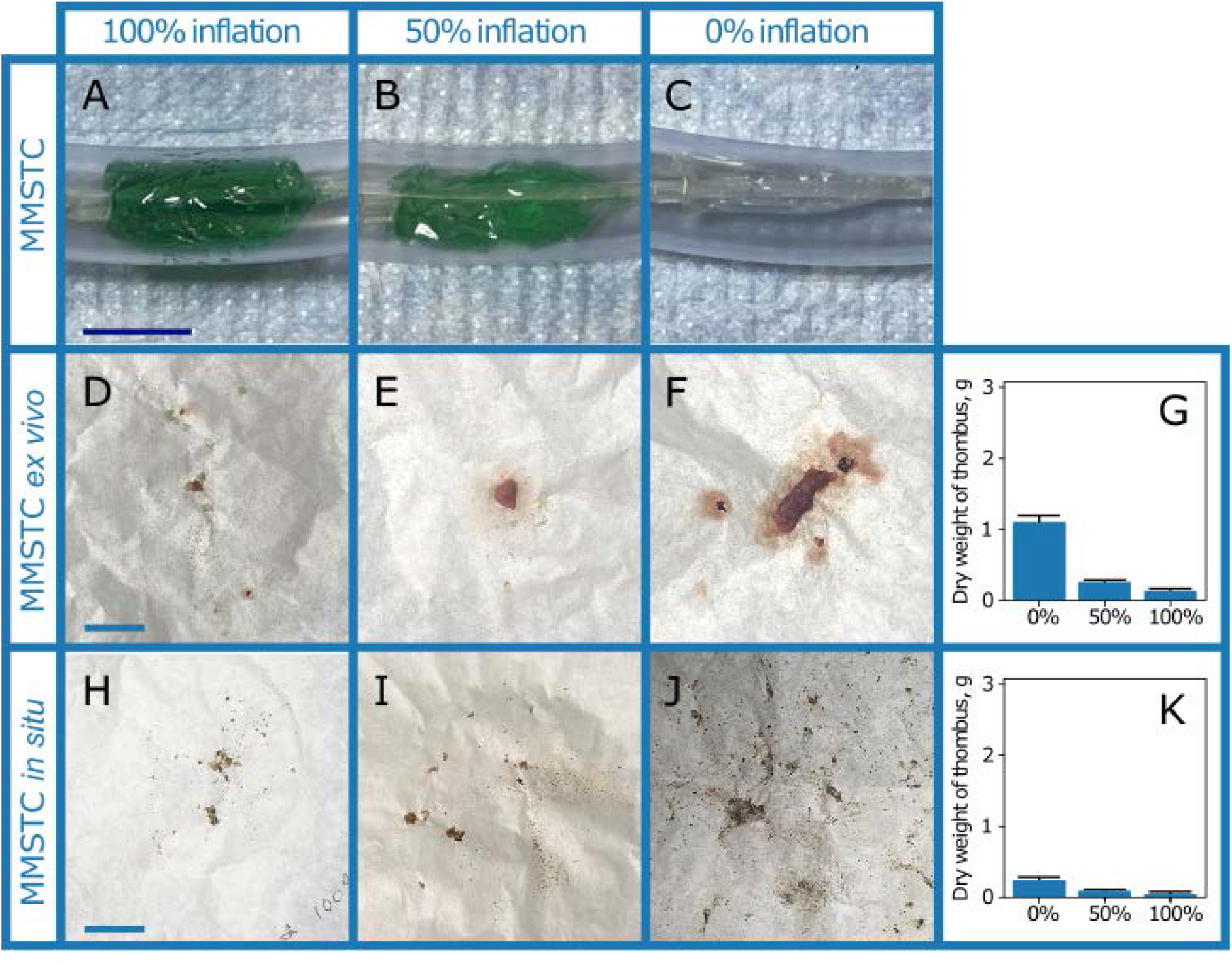
A-C) Multi-modal sequestering thrombectomy catheter (MMSTC) with sequestering distal embolic balloon inflated at 0%, 50%, and 100% in *ex vivo* testing apparatus. D-F) Representative distal emboli collected on filter paper in the laboratory flow system with MMSTC balloons inflated at 0%, 50%, and 100%. G) The balloons caused a significant reduction in distal embolization. H-J) Representative distal emboli collected on filter paper in the laboratory flow system with MMSTC balloons inflated at 0%, 50%, and 100%, in a bovine vena cava segment. K) The balloons again caused a significant reduction in distal embolization.All scale bars: 2 cm.

The average dry weight that converted into distal emboli for the MMSTC was 1.09 ± .10 g at 0% balloon inflation, 0.266± .02 g at 50% balloon inflation, and 0.138± .02 g at 100% balloon inflation (Fig. 6D-G). This indicated that balloon encapsulation of a sequestered treatment zone did indeed significantly reduce distal embolization (*p* < 0.0001, one-way ANOVA). The difference between 100% balloon inflation and 50% balloon was also significant (87.4% relative decrease, p = 0.015).

To assess this issue in a real vein, the procedure was repeated once more, this time in a bovine vena cava segment containing a model thrombus generated as with the thrombi in plastic tubes. With the MMSTC inflation at 0%, 50%, and 100%, the average amount of thrombus embolus was 0.347± .0058 g, 0.190 ±.061 g, and 0.057 ± .010 g, respectively (*p* = 0.039; (Fig. 6H-K)).

When comparing the Penumbra Indigo 5 Fr catheter under identical conditions to the MMSTC *ex vivo* test, 2.69 ± .254 g of distal thrombus emboli. This difference of over an order of magnitude was statistically significant compared to the sequestering MMSTC (*p* = 0.002, (Fig.7 A-C)). The Penumbra Indigo 5 Fr catheter was also compared to the MMSTC under identical *in situ* conditions. The weight of distal emboli collect was found to be 3.16 ± .221 g of thrombus. Under the same conditions, the weight of distal emboli generated from the MMSTC was found to be 0.01 ± .001 g of thrombus, a difference of over an order of magnitude which was statistically significant (*p* < 0.0001, (Fig.7 D-F)).Note that the distal thrombus was larger in the *ex vivo* model than in the *in situ* model because of the flexibility of the vessel wall compared to that of the clear tubing of the flow model.

**Figure 7.**
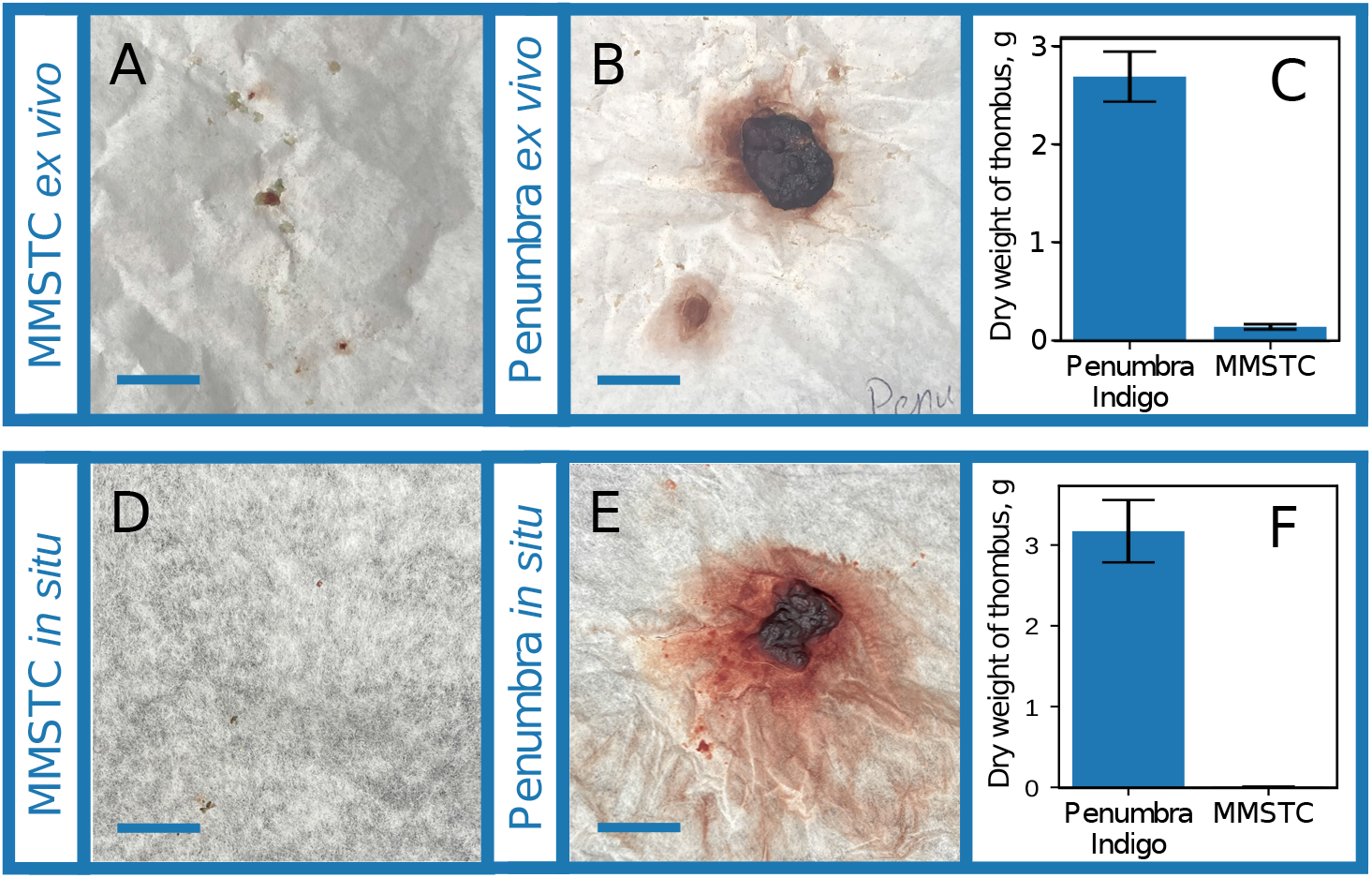
A) Representative distal emboli collected from the MMSTC device *ex vivo*. B) Representative distal emboli collected from the Penumbra Indigo 5 Fr device *ex vivo*. C) The MMSTC device produced more than an order of magnitude less distal embolization than the Penumbra device under identical conditions. D) Representative distal emboli collected from the MMSTC device *in situ*.E) Representative distal emboli collected from the Penumbra Indigo 5 Fr device *in situ*. F) The MMSTC device produced more than an order of magnitude less distal embolization than the Penumbra device under identical conditions. All scale bars: 2 cm.

### Isovolumetric infusion and suction through the MMSTC

To interpret the response of a vena cava segment to isovolumetric infusion, we first studied the response of an idealized thin, hyperelastic tube to suction and positive pressure. The response to a positive pressure is initially compliant locks up at a critical stretch of 1.41. In compression, the tube would have an initially similar compliance. However, the Bryan buckling pressure is exceeded at a pressure of *p/μ* between 0.001 and 0.03 for the range of thicknesses *H/R* considered. Thus, the prediction is that the deformation in response to a suction would be a buckling collapse of the tube (Fig. 8A).

**Figure 8.**
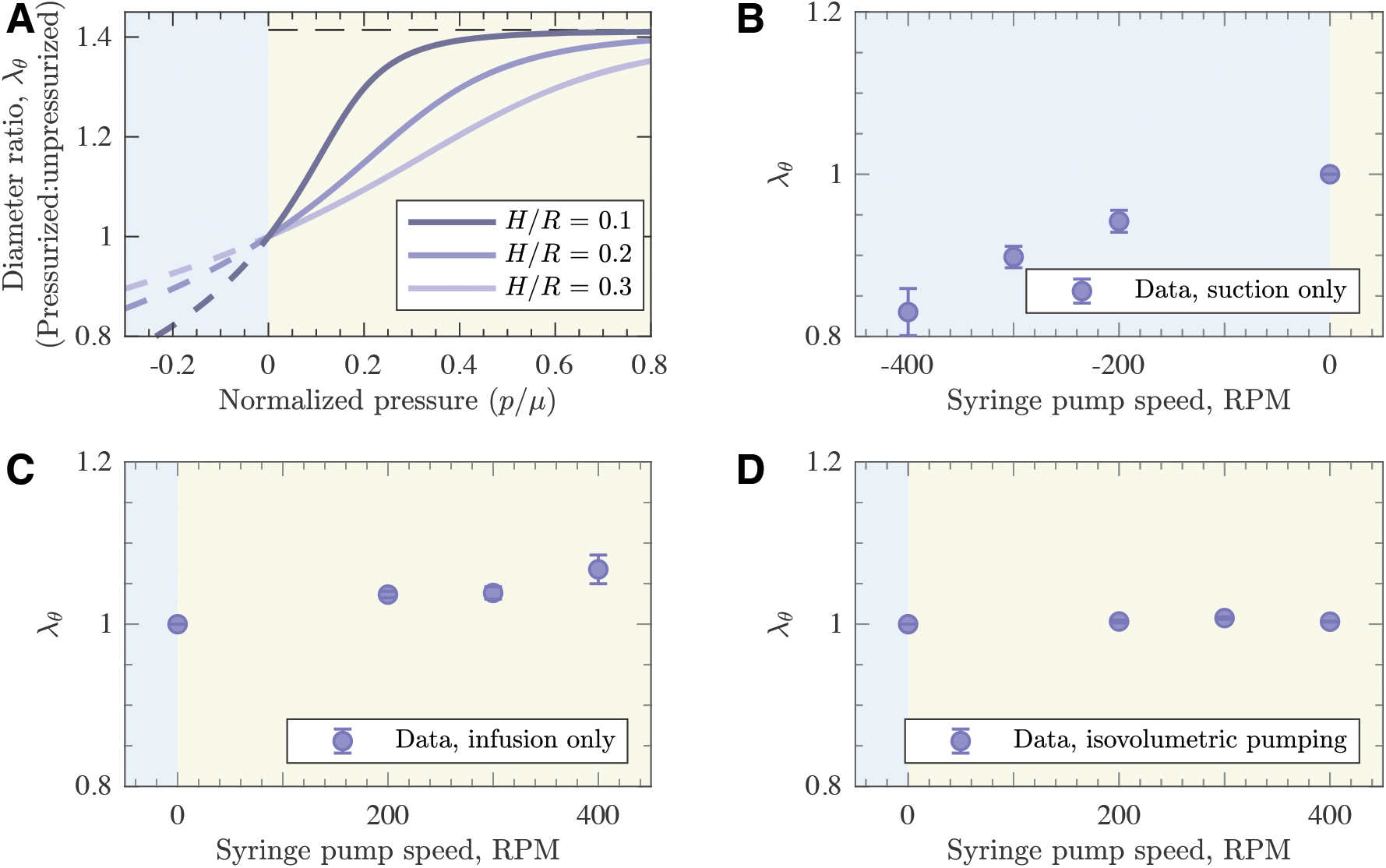
A) A model of the response of a hyperelastic tube to suction and positive pressure loading. The dashed lines represent states that are not achievable due to buckling in compression or material lock-up in tension. B) Change in vena cava diameter (proportional to stretch ratio *λ_θ_*) during suction only using the MMSTC. C) Change in vena cava diameter during infusion only using the MMSTC. D) Change in vena cava diameter during isovolumetric infusion using the MMSTC.

We evaluated these responses in a bovine vena cava specimen *ex vivo*, without a thrombus. The slope of the relationship between pump speed and stretch ratio in the vena cava in suction (Fig. 8B) was much greater than that positive pressure (Fig. 8C). Because pressure increases monotonically with pump speed, this was indicative of buckling in suction as seen in (Fig. 8A).

However, when isovolumetric infusion and suction were performed simultaneously, we observed only a 0.8% increase in vena cava diameter and wall stretch (Fig. 8D). This did not represent a statistically significant change (*p*=0.12). When all three different treatment types where compared to one another we observed that there was a significant difference between treatment type (two-way ANOVA; *p*=0.04).

## Discussion

Distal embolization, vein wall damage, and intraoperative bleeding are important variables when considering procedural safety and outcomes. The multi-modal sequestered therapy may hold promise for treatment of individuals who have suffered a large-volume DVT. The MMSTC prototype demonstrated effective thrombus removal, significant reduction of distal embolization, and a mechanism to decrease vein wall collapse and over-distension during flushing of a defined treatment zone. Furthermore, results with the MMSTC suggest incremental benefits over predicate devices that are already in clinical practice such as the Penumbra Indigo CAT 5 Fr suction thrombectomy catheter. Compared to predicate device, the MMSTC demonstrated a higher thrombectomy efficiency and reduced risk of distal embolization.

The adjustable double-balloon design of the MMSTC was effective at sequestering the thrombus and decreasing distal embolization in an *ex vivo* flow model. Previous studies have used similar thrombosis circuits to evaluate embolus volume, size, and particle counts^10–14^. One study identified that particle counts resulting from distal thrombus emboli can vary in size depending on the type of thrombolyic used^13^. Another study also demonstrated that distal embolization was a common observation with current mechanical thrombectomy devices, highlighting the need for improved next-generation technology that can minimize the risks of distal embolization^10^.

Clinically, the risk of distal thrombus embolization has been widely reported^4, 5, 15–17^. Although it is likely that these events are under-reported, when they do occur, they can lead to catastrophic events including severe hemodynamic compromise, need for invasive measures such as extracorporeal membrane oxygenation (ECMO), or even death. In a recent registry of 234 patients who underwent suction thrombectomy of large volume DVTs in the vena cava and right heart, there was a reported 36 procedure-related complications (15%). Among these complications, one patient with vena cava DVT had intraprocedural distal emboli leading to a massive PE and subsequent death. Of note, transfusion occurred in 59 procedures (25.2%), and in 5% of these patients >3 unites of blood were needed^18^.

The vena cava wall is a highly compliant structure, which is essential for venous capacitance and blood return to the heart depending on an individual’s volume status. With hypovolemia the vein will naturally be constricted, and with hypervolemia the vein wall will be more compliant and distended^19, 20^. However, rapid collapse and over-distension of the vein wall is presumed to cause vein wall micro-trauma, which can lead to denuding of the endothelium, elastin fiber fragmentation, and risk of vein wall thrombosis or internal bleeding^21^.

To date, there is no constitutive relationship or derivation for the vena cava or the venous vasculature in general. Therefore, while the model presented in our study has its limitations (as it relies on constitutive relationships originally defined for rubber biomaterials) it does provide a new method to predict venous wall failure due to excesses suction/infusion pressures through the MMSTC. Simultaneous suction and infusion through the MMSTC led to a 17% change in venous wall diameter (Figure 5), which is far below the theoretical limit of 41% predicting vein wall bursting. There is a paucity of *ex vivo* data that evaluates this process, but our system demonstrated that isovolumetric flow within a vena cava tissue segment can indeed prevent vein wall collapse and over-distension.

Our study results provide a foundation for a sequestered approach to DVT treatment with multi-modal thrombectomy. We anticipate that future pre-clinical studies will help further validate this approach as an effective and safe alterative to existing venous thrombectomy procedures. Future studies will optimize infusion and suction parameters through the MMSTC, utilize proximal and distal balloons that can accommodate a wide range of venous diameters. Additionally, testing of the multi-modal thrombectomy catheter will need to be transitioned to an *in vivo* models to evaluate feasibility and preliminary efficacy of this novel device.

## Author Contributions

M.A.Z. and G.M.G. conceived the design and conduct of the study. U.I. and R.A.R. performed experiments. U.I. and R.A.R. collected the data. U.I., R.A.R., G.M.G., M.A.Z. analyzed and interpreted the data. U.I. wrote the manuscript with major contributions from G.M.G. and M.A.Z.. All authors reviewed and approved the final version of the manuscript. The graphics in Figure 1 was drawn for Caeli Vascular by Matt Holt. Using Solidworks Professional 2019 (https://www.solidworks.com/) J.C drew Figure 2.

## Funding

This work was supported by grants from NIH/NHLBI R41 HL 150963 (M.A.Z), the NSF (CMMI 1548571), and the Skandalaris Center at Washington University in St. Louis.

## Competing Interests

G.M.G. and M.A.Z. have equity in Caeli Vascular, Inc., which specializes in developing treatments for venous disease. U.I. and R.A.R are employed by Caeli Vascular, Inc..

