## Supplement for "Multimodal Thrombectomy Device for Treatment of Acute Deep Venous Thrombosis"

### Mechanics model for vein wall distention and collapse

To study the relationship between a blood vessel volume and pressure, we analyzed the model problem of a hyperelastic cylinder subjected to internal pressure. The vena cava was modeled as a right cylinder of initial wall thickness $H$ and radius $R$. When inflated, the cylinder retains its shape but changes it thickness to $h$ and radius to $r$. The model assumed that the cylinder wall does not rupture and treated the wall as an incompressible, isotropic, Neo-Hookean elastic material. Due to the occlusion of the treatment zone by the distal and proximal balloons, which were connected by a relatively stiff catheter, the length of the cylinder was held constant. Shown in figure 1, the basis of $\left\{\hat{e}_r, \quad \hat{e}_\theta, \quad \hat{e}_z\right\}$,

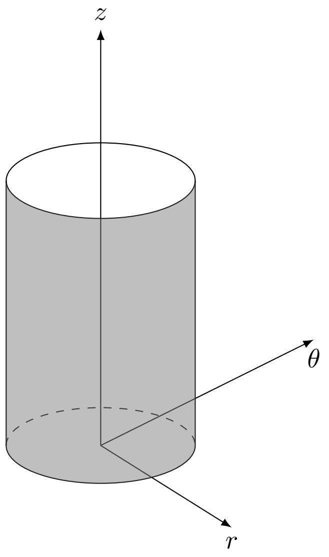


Figure 1: Cylindrical coordinate basis and cylindrical representation of the vena cava.

or the cylindrical coordinate system, will be used.

#### Kinematics

The first step was to relate the changes in the radius of the cylinder to the degree that the material in the cylinder wall stretches. In examining a cross-sectional slice of the hollow cylinder, the three principal stretches ($\lambda_r$, $\lambda_\theta$, and $\lambda_z$) needed to compose the deformation gradient tensor can be written, where a principal stretch is defined as the ratio of the deformed length of a piece of material in one of the three coordinate directions ($r$, $\theta$, or $z$) to the initial length of that piece of material in that coordinate direction.

$\lambda_r$, the stretch ratio in the radial direction, is the ratio of initial to final wall thickness:

$$\lambda_r = \frac{h}{H}. \tag{1}$$

$\lambda_\theta$, the stretch ratio in the "hoop" (circumferential) direction, is the ratio of final to initial circumference:

$$\lambda_\theta = \frac{2\pi r}{2\pi R} = \frac{r}{R} \tag{2}$$

Finally, the stretch $\lambda_z$ along the axis of the cylinder is unity because the length of the treatment zone is fixed by the connection between the proximal and distal balloons, the final length $l$ and initial length $L$ are identical, so that:

$$\lambda_z = \frac{l}{L} = 1 \tag{3}$$

To enforce the constraint that the wall of the vena cava as incompressible, we note that the initial and final volume are $V = 2\pi RLH = 2\pi rlh$, so that

$$\frac{2\pi rlh}{2\pi RLH} = \lambda_r\lambda_\theta = 1, \tag{4}$$

or $\lambda_r = \lambda_\theta^{-1}$.

**Stresses**

The next step is to examine the three stresses in the wall of the cylinder: $\sigma_r$, $\sigma_\theta$, and $\sigma_z$. Because the thickness of the cylinder is significantly smaller than its radius, $\sigma_r$ can be neglected relative to the other terms so that the stress state is approximately two-dimensional. These stresses can be calculated from the "Law of Laplace" by treating the cylinder as a membrane in equilibrium with an internal pressure, $p$, that arises from fluid inside the vena cava or the thrombus itself, and acts uniformly on the internal lumen and both ends of the vena cava. Considering the equilibrium of a cross-section of the vena cava, the a stress $\sigma_\theta$, acting over the circumferential cross-sectional area $2\pi rh$, must balance the internal pressure acting over an area $\pi r^2$, so that $2\pi rh\sigma_z = \pi r^2 p$, or:

$$\sigma_z = \frac{pr}{2h} \tag{5}$$

A second force balance can be achieved by examining the cylinder if it were cut in half lengthwise. In this case, $p$ acts over an area $2rl$, and is counteracted by $\sigma_\theta$ which acts on the membrane walls of area $2hl$, so that $\sigma_\theta hl = prl$, or:

$$\sigma_\theta = \frac{pr}{h} \tag{6}$$

#### Constitutive Behavior

Finally, the constitutive relation that relates stress to stretch of the vena cava was considered. The vena cava was modeled as isotropic and neo-Hookean, so that:

$$\sigma_\theta = 2\mu J^{-5/3}(\lambda_\theta^2 - \lambda_r^2) \tag{7}$$

$$\sigma_z = 2\mu J^{-5/3}(\lambda_z^2 - \lambda_r^2) = 2\mu J^{-5/3}(1 - \lambda_r^2) \tag{8}$$

where $J = \lambda_r \lambda_\theta \lambda_z = \lambda_r \lambda_\theta$. Substituting:

$$\frac{pr}{h} = 2\mu J^{-5/3}(\lambda_\theta^2 - \lambda_r^2) \tag{9}$$

$$\frac{pr}{2h} = 2\mu J^{-5/3}(1 - \lambda_r^2) \tag{10}$$

Solving:

$$1 - \lambda_r^2 = \lambda_\theta^2 - 1 \tag{11}$$

Substituting once more:

$$\frac{pr}{h} = p\frac{\lambda_\theta R}{\lambda_r H} = 2\mu J^{-5/3}(2\lambda_\theta^2 - 2) \tag{12}$$

$$p = 2\mu \frac{\lambda_r H}{\lambda_\theta R} \frac{(2\lambda_\theta^2 - 2)}{(\lambda_r \lambda_\theta)^{5/3}} \tag{13}$$

$$= 4\mu \frac{H}{R} \frac{(\lambda_\theta^2 - 1)}{(\lambda_r^2 \lambda_\theta^8)^{1/3}} \tag{14}$$

$$= 4\mu \frac{H}{R} \frac{(\lambda_\theta^2 - 1)}{(2\lambda_\theta^8 - \lambda_\theta^{10})^{1/3}} \tag{15}$$
